# Electrophysiological indices of individual differences in adult language learning

**DOI:** 10.1101/2022.06.07.495229

**Authors:** Halima Nalaye, Zachariah R. Cross, Matthias Schlesewsky, Ina Bornkessel-Schlesewsky

**Affiliations:** Cognitive Neuroscience Laboratory – Australian Research Centre for Interactive and Virtual Environments, University of South Australia, Adelaide, South Australia, Australia

**Keywords:** individual alpha frequency, language learning, modified miniature language, N400, EEG

## Abstract

Individual differences in second language (L2) learning can offer insights into the neurobiological bases of learning aptitude. One neurophysiological marker of inter-individual differences in cognition is the individual alpha frequency (IAF), a trait-like measure correlated with cognition. Further, the N400 is an electrophysiological marker indexing stimulus irregularity and has been used to study L2 learning; however, its relationship with IAF and L2 learning remains unknown. To examine the relation between IAF and L2 learning (indexed by N400 amplitude), we report data from a modified miniature language learning study. After a vocabulary learning period, participants (*N* = 38, M_age_ = 25.3, SD = 7.13) judged the grammaticality of classifier-noun pairs, with mixed-effects modelling revealing lower IAF individuals were better than higher IAF individuals at grammaticality judgements. N400 amplitude also reduced across the experiment in low relative to high IAF individuals, indicating the relationship between IAF and language learning is more complex than initially postulated.

## 1. Introduction

Learning a second language (L2) in adulthood is considerably more challenging than in childhood, with only a small portion of learners meeting native-like proficiency (Prat et al., 2016; Zafar & Meenakshi, 2012). Language acquisition is a multi-faceted process, involving the rapid formation of syntactic and semantic knowledge and the ability to make logical inferences in one’s target L2 (Kepinska et al., 2018). However, compared to children’s general success in language learning, adult L2 learners show deficits in phonological, syntactic, and semantic knowledge acquisition in their target language (Lenneberg, 1967; Mueller, 2006). Several factors are posited to explain this phenomenon, particularly the age of acquisition of one’s target L2 (Flege et al., 1999).

It is often argued that there is a critical period for language acquisition that limits the ability for adult learners to achieve native-like proficiency, a viewpoint first endorsed by Penfield & Roberts (1959). However, research demonstrates that a portion of adult L2 learners become so skilled at their target language that they are indistinguishable from native speakers (e.g., Birdsong, 1992; White & Genessee, 1996). This point towards a need for further research into the factors underlying inter-individual differences in L2 learning, possibly providing further insight into the neural underpinnings of language learning and other forms of complex information processing (Kepinska et al., 2017). There are several methodologies with which L2 learning can be studied experimentally, two of the most common being artificial grammar learning and modified miniature language paradigms.

### 1.1. Simulating the structural regularities of language using artificial grammar learning

Artificial grammar learning (AGL) paradigms consist of specific rules used to generate strings of grammatical sequences presented to participants during an exposure or learning phase (Friederici et al., 2002; Kepinska et al., 2018; Pothos, 2007). During testing, novel stimuli that either follow or violate these rules are presented to test participants’ knowledge (Westphal-Fitch et al., 2018). Although participants are often not made explicitly aware of these rules, previous research has shown that participants can, albeit with varying success, classify presented stimuli of varying complexity as grammatical or ungrammatical (Knowlton & Squire, 1994). While the complex nature of natural language makes it difficult for researchers to control for variables such as prior learning (Folia et al., 2010), AGL paradigms allow for the analysis of the acquisition of specific structural aspects of language (e.g., Gómez & Gerken, 2000; Petersson et al., 2012).

Despite the widespread use of AGL paradigms to study language learning, AGL paradigms cannot be easily generalised to natural language learning (Mueller, 2006). AGL studies often only require learning artificially constructed rules and, unlike natural languages, often do not contain meaning (Fitch & Friederici, 2012; Rebuschat & Williams., 2012). An alternative paradigm for studying higher-order L2 learning, which goes beyond syntax alone, are modified miniature languages (e.g., Cross et al., 2020a; Mueller, 2006).

### 1.2. Reintroducing meaning into artificial language: Modified miniature languages

Modified miniature languages (MMLs), like AGLs, are a psycholinguistic paradigm that enables precise control over stimuli, allowing isolation of qualities of interest such as particular grammatical rules (e.g., syntactic rules; Mueller, 2006). However, unlike AGLs, MMLs are based on natural languages (e.g., Cross et al., 2020a; Mueller, 2006). The construction of MMLs thus lends them well to studying L2 acquisition, as their basis on pre-existing languages mimics language in an arguably more ecologically valid way than AGLs (Mueller et al., 2005).

Despite the body of research into L2 learning using experimental methods such as MMLs (Cross et al., 2020a; Mueller, 2006), relatively little is known regarding how individual differences in neurophysiology influence the speed at which non-native speakers learn an L2 (cf. Cross et al., 2022). Most research has focused on the role cognitive mechanisms play in L2 learning, including implicit learning (Denhovska & Serratrice, 2017), memory (Morgan-Short et al., 2014), and individual factors such as motivation (for a review, see Dörnyei, 2005). While these factors play a role in language learning, the influence of inter-individual differences in neurophysiology has not been extensively studied in this domain.

One way to index such differences is by analysing resting-state-derived brain activity (Chai et al., 2016), which reflects the global functional capacity of the brain (Fox et al., 2007). One such example of resting-state brain activity associated with a range of cognitive domains is the individual alpha frequency (Grandy et al., 2013b).

### 1.3. Individual alpha frequency as a neurophysiological proxy for individual differences in cognition

Since the discovery of the human EEG by Berger (1929), the relationship between neurophysiological activity within the alpha range (∼7 – 13 Hz) and cognitive ability has been widely studied (Surwillo, 1963; Klimesch, 1999; Klimesch et al., 1993). Individual alpha frequency (IAF) is associated with memory (Lebedev, 1994), language (Bornkessel et al., 2004; Bornkessel-Schlesewsky et al., 2015), attention (Angelakis et al., 2004, Klimesch et al., 1993), general cognitive ability (g-factor intelligence; Grandy et al., 2013a, Zakharov et al., 2020), and has recently been shown to modulate sleep-related memory consolidation (Chatburn et al., 2021; Cross et al., 2020b). Specifically, a growing body of work shows that IAF is a more sensitive measure of individual cognitive performance than behavioural measures alone in studying language comprehension (Bornkessel et al., 2004; Bornkessel-Schlesewsky et al., 2015; Kurthen et al., 2020), as well as in naturalistic paradigms of cognition (Dziego et al., 2022)

The functional significance of IAF as a robust marker of information processing makes it a potentially informative metric for examining individual differences in L2 learning, as it is advantaged by its stable, trait-like characteristics (Posthuma et al., 2001) and test-retest reliability in samples of healthy individuals (Grandy et al., 2013b). However, the relationship between IAF and L2 learning aptitude has not been studied to date. One such measure to quantify language learning at an electrophysiological level is the N400 event-related potential (ERP), which has been used extensively in previous studies of non-native speaker L2 acquisition as a marker of learning is the N400.

### 1.4. The N400 as an electrophysiological marker of L2 learning

Language learning is a rapid process involving the extraction and generalisation of complex rules into long-term memory (Cross et al., 2018, 2021; Mickan et al., 2019; Osterhout et al., 1997). Previous studies show that the processing of ecologically valid artificial languages, such as MMLs, engender similar electrophysiological correlates to those found in natural language processing (Mueller et al., 2007). A negative deflection peaking around 400ms post-stimulus onset with a centro-parietal distribution, the amplitude of the N400 marks how anomalous the occurrence of a presented stimulus is (Kutas & Federmeier, 2011; Kutas & Hillyard, 1980).

The amplitude of the N400 is linked to the degree of stimulus predictability (Frank et al., 2015, Lau et al., 2008), with predictable words in an appropriate context resulting in reduced N400 amplitude. From this perspective, the amplitude of the N400 can be used as a proxy for the predictability of sensory input, with stronger amplitude indicating a higher degree of ‘unexpectedness’ (Terporten et al., 2018). Further, as N400 effects are often seen in the absence of behavioural differences, the N400 may serve as a more sensitive measure of language learning. For example, McLaughlin et al. (2004) found that adult learners showed subtle N400 effects corresponding to language learning after 14 hours of instruction, effects which were not detected with behavioural assessment alone.

The role electrophysiological markers such as the N400 play in L2 acquisition has been previously studied. However, previous research has not examined N400 amplitude in conjunction with inter-individual differences in electrophysiology (as quantified by IAF) as a predictor of L2 learning aptitude. As there is extensive inter-individual variability in language learning ability, and given that IAF reflects differences in information processing, it may help to explain more variance in language learning than previously used methods such as standard behavioural measures of learning alone. One method of studying this relationship is examining L2 learners’ ability to generalise grammatical features novel to them into their existing knowledge (with this ability quantified by N400 amplitude).

### 1.5. Mandarin Chinese and classifiers

By studying how adult L2 learners integrate implicit novel grammatical rules into associative memory using indices such as the N400, researchers can infer the neural processes governing language acquisition. One grammatical feature novel to monolingual native English speakers is classifiers. Classifiers are an additional type of morpheme that groups objects numerically and are found in East Asian languages (e.g., Mandarin Chinese; hereafter referred to as Mandarin; Gao & Malt, 2009). Classifiers provide linguistic categorisation of the world by sorting objects into classes (Zhang, 2007). For example, whereas in English one would say *a rope*, a Mandarin speaker would say *yi tiao shengzi*, with *yi* being the numeral *one*, *shengzi* meaning *rope*, and *tiao* being a classifier denoting a *long thing* – translated as *one long-thing rope* (for more information, see Gao & Malt, 2009). The N400 as an electrophysiological index of L2 learning has been examined in previous studies of classifier learning in Mandarin (Chou et al., 2014; Hsu et al., 2014; Kwon et al., 2017; Qian & Garnsey, 2016) and other East Asian languages such as Japanese (Kanero et al., 2015; Muller et al., 2005) and Korean (Jin, 2018), where classifier-noun pair violations elicited an N400.

In all East-Asian languages where classifiers are a grammatical feature such as Japanese, classifier choice must be congruent with the noun that the classifier is modifying (Qian & Garnsey, 2016). Associative memory – memories regarding the relationship between items (Dennis et al., 2015) – has been linked with the ability to discern appropriate classifier-noun choice (Cross et al., 2020a). As a higher IAF is linked with improved memory performance across domains (Klimesch, 1999), possessing a higher IAF may influence the ease of making an appropriate classifier-noun choice. The functional role that IAF plays in memory may therefore manifest itself in an individual’s N400 response, with a stronger amplitude indicating a greater degree of associative memory strength. This may consequently result in a greater ability for L2 learners to implicitly learn and generalise the rules governing appropriate classifier-noun pairing in sentence processing.

### 1.6. The current study

The present study examined individual differences in L2 learning. To this end, behavioural and electrophysiological data were reanalysed from Cross et al. (2021), which employed an MML paradigm derived from Mandarin, hereafter referred to as ‘Mini-Pinyin’. The primary aim was to examine the relationship between IAF and learning novel grammatical rules. Classifier-noun pairing grammaticality judgement accuracy and N400 amplitude served as measures of learning, on which effects of IAF could be examined. Single-trial N400 amplitude was the primary outcome measure of online classifier processing, given its demonstrated sensitivity to violations in the early stages of language learning.

It was hypothesised that (H^1^) response accuracy would increase across the experiment due to an increased sensitivity to syntactic violations as a function of learning; (H^2^) response accuracy would be greater in those with a higher IAF compared to those with a lower IAF; (H^3^) the amplitude of the N400 in response to classifier violations would increase throughout the experiment, due to an increased sensitivity to syntactic violations as more learning and integration of grammatical knowledge occurs; (H^4^) the N400 effect will be more pronounced in those with a higher IAF compared to those with a lower IAF, and; (H^5a^) the N400 will decrease across the course of the experiment for high relative to low IAF individuals, and (H^5b^) this difference will be larger for classifier-noun violations at the first relative to second noun phrase position.

## 2. Methods

### 2.1. Participants

Data obtained from 38 participants were used for analysis in this study (mean age = 25.3, *SD* = 7.13). Key inclusion criteria required individuals to be monolingual right-handed (as measured by the Flinders Handedness Survey; FLANDERS; Nicholls et al., 2013) native English speakers with no prior exposure to Mandarin, as well as meeting general inclusion criteria relevant to EEG studies. Informed consent was gained from all participants, with each participant receiving a $120 honorarium. Data collection was conducted at the University of South Australia’s Cognitive Neuroscience Laboratory, with ethics approval obtained from the University of South Australia Human Research Ethics Committee (approval I.D. 0000032556).

### 2.2. Measures

#### 2.2.1. Modified miniature language paradigm

The modified miniature language paradigm used in this study was Mini Pinyin, which is based on Mandarin Chinese (Cross et al., 2020a). Mini Pinyin is comprised of 16 transitive verbs, 25 nouns, two coverbs, and four classifiers. Each category of noun is associated with a specific classifier that precedes each noun in a sentence (see Table 1). For a complete description of the structure of Mini Pinyin, see Cross et al. (2020a).

**Table 1.** Noun Category and Corresponding Classifier in Mini Pinyin.

| Noun Category | Classifier |
| --- | --- |
| Animal | zhi |
| Human | ge |
| Object (big) | da |
| Object (small) | xi |

In the learning phase of the experiment, participants were provided with a vocabulary booklet consisting of the 25 nouns Mini-Pinyin is comprised of. This booklet solely consisted of the nouns, with all other grammatical features being learned during the sentence learning phase of the experimental session. After an independent learning period which lasted a minimum of three and a maximum of seven days, participants returned to the lab to complete the experiment.

Regarding the syntactic constraints of sentence construction in Mini Pinyin, there are two types of manipulations surrounding word order rules, and classifier-noun pairs, with the latter examined in this study. In grammatical sentences, classifiers are consistent with their associated noun-pair (e.g., human [ge] paired with the word for pirate [haido]). By contrast, ungrammatical sentences violate this rule by pairing classifiers with an inappropriate noun (e.g., human [ge] with the word for horse [junma]; see Table 2 for examples of grammatical and ungrammatical sentence constructions). These manipulations allow for the measurement of individual responses to classifier-noun pair violations at either the first or second noun phrase position. Two lists of sentence stimuli were created, with 144 being grammatical and 48 being ungrammatical classifier-noun violations.

**Table 2.**
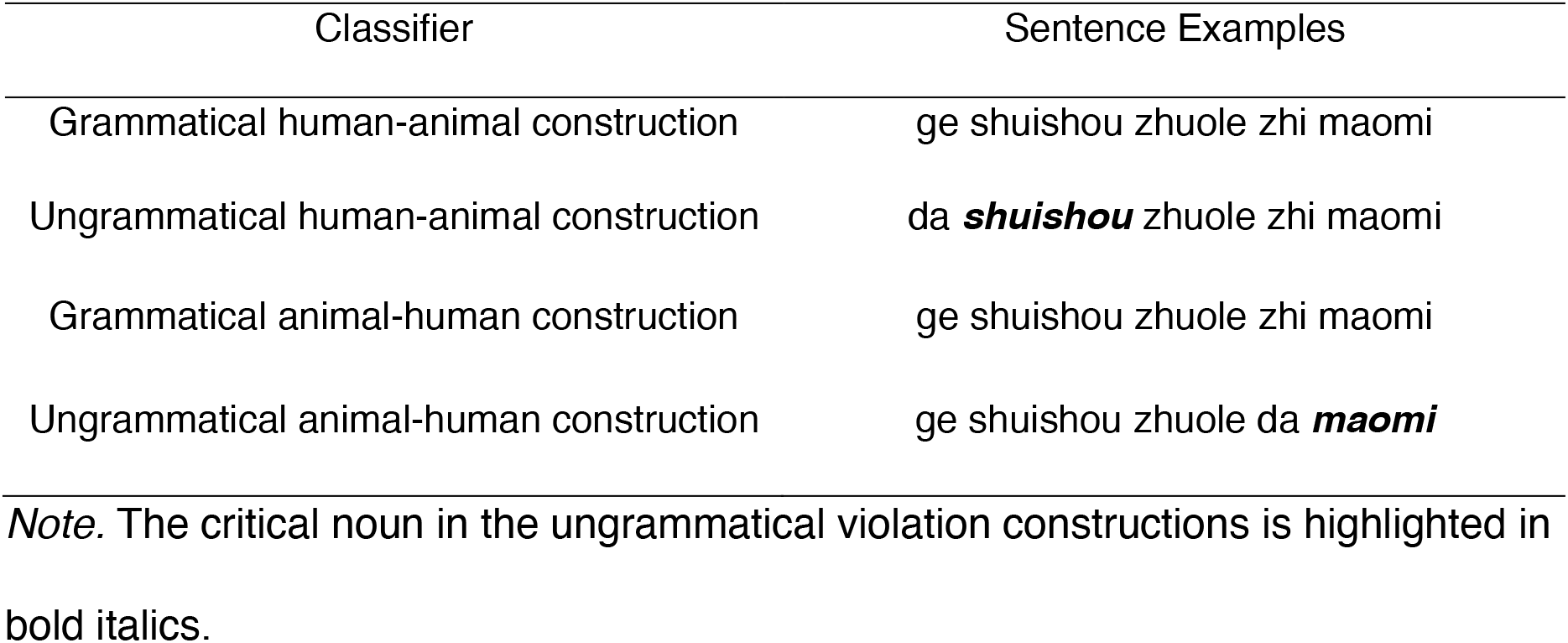
Example of Grammatical and Ungrammatical Classifier-Noun Pair Sentence Constructions in Mini-Pinyin.

| Classifier | Sentence Examples |
| --- | --- |
| Grammatical human-animal construction | ge shuishou zhuole zhi maomi |
| Ungrammatical human-animal construction | da ***shuishou*** zhuole zhi maomi |
| Grammatical animal-human construction | ge shuishou zhuole zhi maomi |
| Ungrammatical animal-human construction | ge shuishou zhuole da ***maomi*** |

*Note.* The critical noun in the ungrammatical violation constructions is highlighted in bold italics.

#### 2.2.2. Electroencephalography

EEG was recorded during quiet resting wakefulness and during the sentence judgement task using a BrainCap 32-channel standard cap with Ag/AgCl electrodes (BrainProducts GmbH, Gilching, Germany), mounted according to the 10-20 electrode placement system. The reference was located at FCz and ground at AFz. EEG signals were re-referenced to linked mastoids offline. Electrooculogram (EOG) was recorded to control for artefacts resulting from eye movements via electrodes at the outer canthus of each eye (horizontal EOG) and above and below the participants’ left eye (vertical EOG). The EEG was amplified using a BrainAmp DC amplifier (BrainProducts GmbH, Gilching, Germany), with an initial band-pass filter of DC – 250 Hz and a sampling rate of 1000 Hz. Impedances were kept below 10k Ω.

### 2.3. Procedure

#### 2.3.1. Vocabulary test

Participants first undertook a computerised vocabulary test, where they translated the learned nouns from Mini-Pinyin to English using a keyboard. Each trial began with a 600ms fixation cross, followed by a visual presentation of the noun form (e.g., a picture of a dog), with 20 seconds to respond. Participants were required to score >90%, with those scoring lower unable to participate in the main experimental session. All participants included in the current analysis scored above 90%.

#### 2.3.2. Sentence learning task

The next phase involved sentence learning, with sentence and picture stimuli presented via OpenSesame (Mathôt et al., 2012), which depicted events occurring between two entities. Although participants were aware that they would complete a sentence learning task, no explicit description of grammatical rules was provided. A fixation cross was first presented for 1000ms, followed by a picture depicting the occurring event for 5000ms. A sentence describing the event in the picture was then presented on a word-by-word basis, with each word presented for 700ms and followed by a 200ms inter-stimulus interval (ISI). This procedure is detailed in Figure 1 and occurred for all 128 sentence-picture combinations. Including three self-paced breaks, the task took approximately 40 minutes to complete.

**Figure 1.**
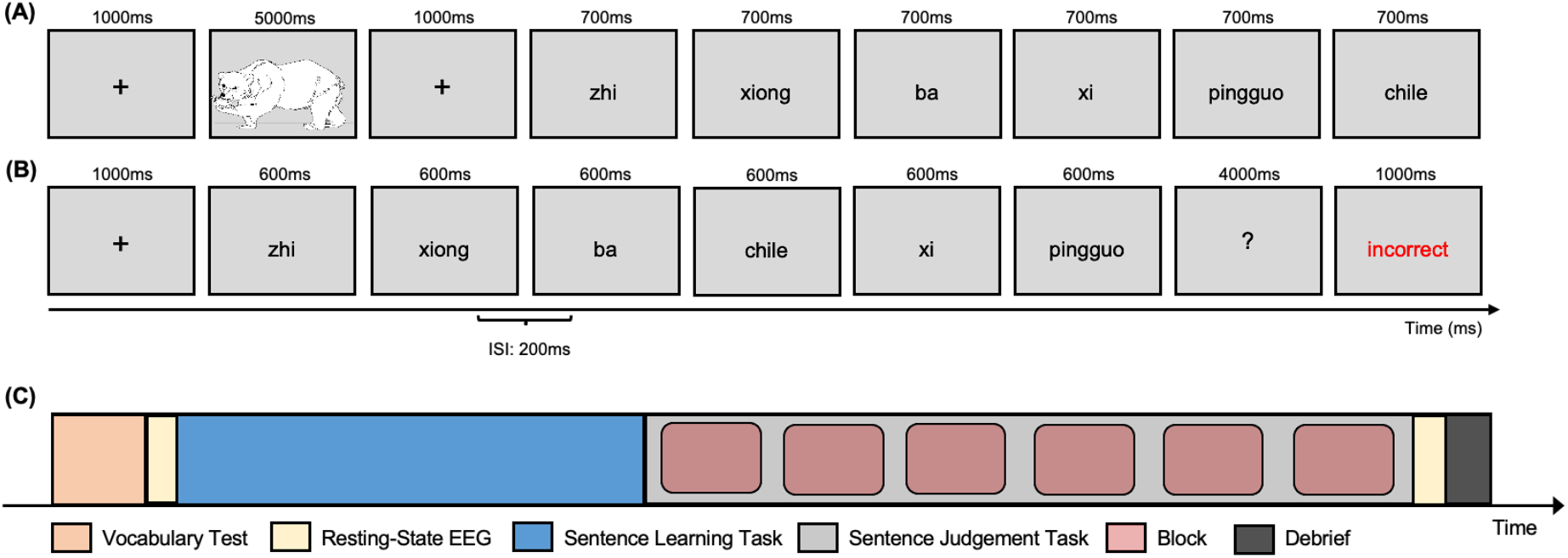
(A) Schematic representation of the sentence learning task. (B) Schematic representation of the sentence judgement task. (C) Main experimental protocol.

#### 2.3.3. Sentence judgement task

Immediately after the sentence learning task, participants completed a sentence judgement task, illustrated in Figure 1. 144 grammatical and 144 ungrammatical (48 of which were analysed here) novel sentences without pictures were presented on a word-by-word basis via rapid serial visual presentation, with each word presented for 600ms and an ISI of 200ms. Feedback was provided after each response to facilitate learning. Stimulus presentation was randomised so that sentences following the same grammatical structure were not judged twice in a row. Participants were instructed to pay attention and to judge the grammaticality of presented sentences (yes/no response) by keyboard presses. To facilitate judgement, a cue of a question mark appeared in the middle of the screen for 4000ms after the presentation of the last word, followed by feedback on the participant’s response, delivered by immediately presenting the word ‘correct’ presented in green text or ‘incorrect’ in red text for 1000ms. The sentence judgement task consisted of six learning blocks, with self-paced breaks between each block.

### 2.4. Main experimental protocol

Participants were given a paired picture-word vocabulary booklet containing the 25 Mini-Pinyin nouns, which they were required to learn at home to ensure a basic understanding of the language. After approximately one week, participants were invited to the Cognitive Neuroscience Laboratory for the main experimental session. Participants were fitted with an EEG cap, with eyes-open and eyes-closed resting-state EEG measured to estimate IAF. Participants then undertook the vocabulary test, followed by the sentence learning and sentence judgement tasks. The complete experimental protocol is detailed in Figure 1.

## 3. Data Analysis

### 3.1. EEG pre-processing

EEG data were pre-processed and analysed using MNE-Python (Gramfort et al., 2014). The raw EEG signal was first filtered with a digital phase true-finite impulse response (FIR) band-pass filter (0.1-30 Hz). EOG artefacts were then identified and removed using the ‘mne.pre-processing.find_eog_events’ function and Independent Component Analysis (ICA; FastICA). Epochs with peak-to-peak amplitude over 75 microvolts or under 5 microvolts were also excluded. Trial-by-trial pre-stimulus amplitude (−200ms to 0ms) was extracted from each epoch and used as a covariate in statistical analyses, as recommended by Alday (2019).

### 3.2. N400 amplitude

Trial-by-trial EEG amplitude in the time window of 400-600ms relative to target word onset was calculated to capture the N400 and was estimated from central and parietal electrodes, given the prominence of N400 activity over these regions (Kutas & Hillyard, 1980). As classifiers precede each noun-pair in a sentence, the point of violation measured was, therefore, the noun. For example, if the human (*ge*) classifier were paired with the word *dog*, the point of violation would be *dog*.

### 3.3. IAF estimation

IAF estimates were obtained using the ‘philistine.mne.savgol_iaf’ function implemented in MNE-Python based on the procedure detailed in Corcoran et al. (2018). This routine calculates the centre of gravity and peak alpha frequency (PAF), with PAF used in the current analyses. Participants’ PAF were estimated from occipital-parietal electrodes, given the prominence of alpha activity over these regions during rest-state eyes closed recordings (Salenius et al., 1995).

### 3.4. Behavioural response accuracy

Grammaticality judgments were calculated on a trial-by-trial basis, determining whether participants correctly identified grammatical and ungrammatical sentences.

### 3.5. Statistical analysis

Statistical analyses were conducted using *R* v.4.0.2 (R Core Team, 2020) with packages *tidyverse* v.1.3.0 (Wickham et al., 2019), *car* v.3.0.8 (Fox & Weisberg, 2019), *effects* v.4.1.4 (Fox & Weisberg, 2019) and *lme4* v.1.1.23 (Bates et al., 2015). Plots were generated using *ggplot2* v.3.3.0 (Wickham, 2016). *lmerOut* v.0.5 (Alday, 2019) was used to produce model output tables, while *ggeffects* v.1.0.2 (Lüdecke, 2018) was used to extract modelled effects for visualisation.

Data were analysed using (generalised) linear mixed-effects models fit by restricted maximum likelihood (REML) estimates. The behavioural model took the following form:

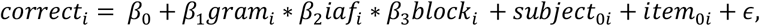

Here, *correct* refers to the binomial response of correct (1) or incorrect (0) during the sentence judgement task, *gram* encodes the grammaticality of the sentence (grammatical, ungrammatical), *iaf* is individual alpha frequency estimates, and *block* is experimental block (1 – 6) from the sentence judgement task. Subject and item were also modelled as random effects on the intercept. The EEG model took the following form:

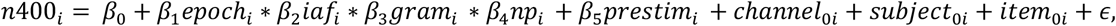

Here, *n400* refers to single-trial N400 amplitude estimates, *epoch* encodes information relating to the timing of each stimulus, with the first epoch indicating the start of the experiment. *iaf* is the individual alpha frequency, and *gram* encodes the grammaticality of the sentence (grammatical, ungrammatical), while *np* refers to the classifier-noun phrase position (first versus second noun phrase). Electrode, subject and item were modelled as random effects on the intercept, while pre-stimulus amplitude was modelled as a covariate (Alday, 2019). In both models, asterisks denote interaction terms, including all subordinate main effects; pluses denote additive terms.

Type II Wald χ2-tests from the *car* package (Fox, 2012) were used to provide *p*-value estimates, while categorical factors were sum-to-zero contrast coded, such that factor level estimates were compared to the grand-mean (Schad et al., 2020); however, block was specified as an ordered factor. Further, an 83% confidence interval (CI) threshold was used given that this approach corresponds to the 5% significance level with non-overlapping estimates (Austin & Hux, 2002; MacGregor-Fors & Payton, 2013). In the visualisation of effects, non-overlapping CIs indicate a significant difference at *p* < .05.

## 4. Results

### 4.1. Descriptive statistics

Overall, participants performed better at identifying grammatical sentences on average (M_correct_ = 65.93%, *SD* = 19.3%) than rejecting ungrammatical sentences (M_correct_ = 50.00%, *SD* = 23.69%). See Figure 2 for a summary of behavioural performance and IAF estimates.

**Figure 2.**
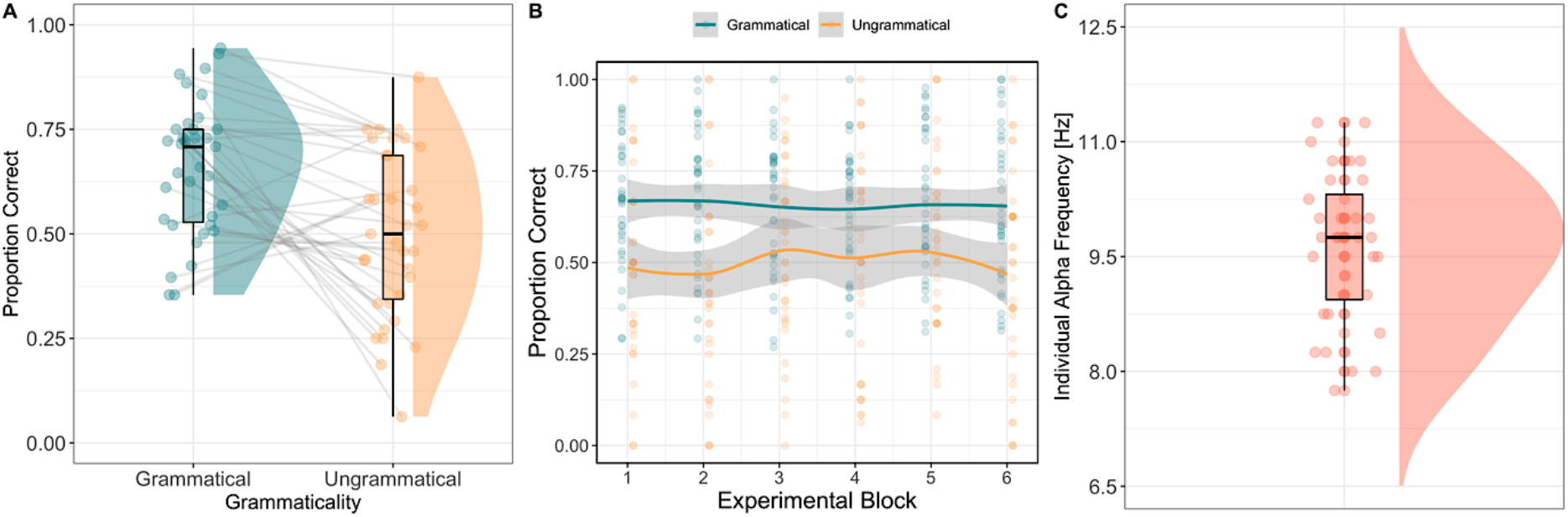
(A) Proportion of correct responses for grammatical and ungrammatical sentences. Data points represent individual participants, while the lines link within-participant performance between grammatical and ungrammatical sentences. (B) Proportion of correct responses for grammatical and ungrammatical sentences across experimental blocks (1 – 6). Data points represent individual participants, while the solid blue line represents grammatical sentences, and the solid yellow line represents ungrammatical sentences. The shaded region indicates the standard error of the mean. (C) distribution of IAF estimates (Hz). Data points represent individual participants.

### 4.2. Modelling behavioural accuracy across time, grammaticality and iaf

To address the first hypothesis that response accuracy would increase over the experiment, a generalised linear mixed-effects model was constructed to examine the interaction between sentence grammaticality and experimental time (i.e., block). A significant main effect of Grammaticality was found (χ2(1) = 17.85, *p* <.001); however, the main effect of Block was non-significant (χ2(5) = 3.75, *p* = .59). Additionally, there was no significant interaction between Grammaticality and Block (χ2(5) = 10.48, *p* = 0.06). This indicates that although there were changes in accuracy across the experiment (Figure 2B), there was no significant increase in participants correctly identifying the grammaticality of sentences over the experiment.

A generalised linear mixed model was constructed to address the second hypothesis that response accuracy will show a larger increase over the course of the experiment in higher IAF compared to lower IAF individuals. However, this initial model did not converge. Block was subsequently removed from the model, given that there was no significant main effect of Block in the previous model. This subsequent model examined the relationship between grammaticality and IAF in predicting the probability of a correct response.

A significant main effect of Grammaticality (χ2(1) = 95.17, *p* < .001) was found, where grammatical sentences had higher response accuracy than ungrammatical sentences. A significant effect of IAF (χ2(1) = 4.44, *p* = .03; see Figure 5A) was found, where those with a lower IAF had a greater accuracy rate than those with a higher IAF. However, the interaction between Grammaticality and IAF was nonsignificant (χ2(1) = 0.50, *p* = .48), indicating that the grammaticality of the sentence and IAF do not interact to influence the probability of a correct response. Interestingly, the relationship observed is in the opposite direction to what was hypothesised, namely that those with a higher IAF would perform better in comparison to those with a lower IAF (for summaries of all models, see the supplementary material).

**Figure 3.**
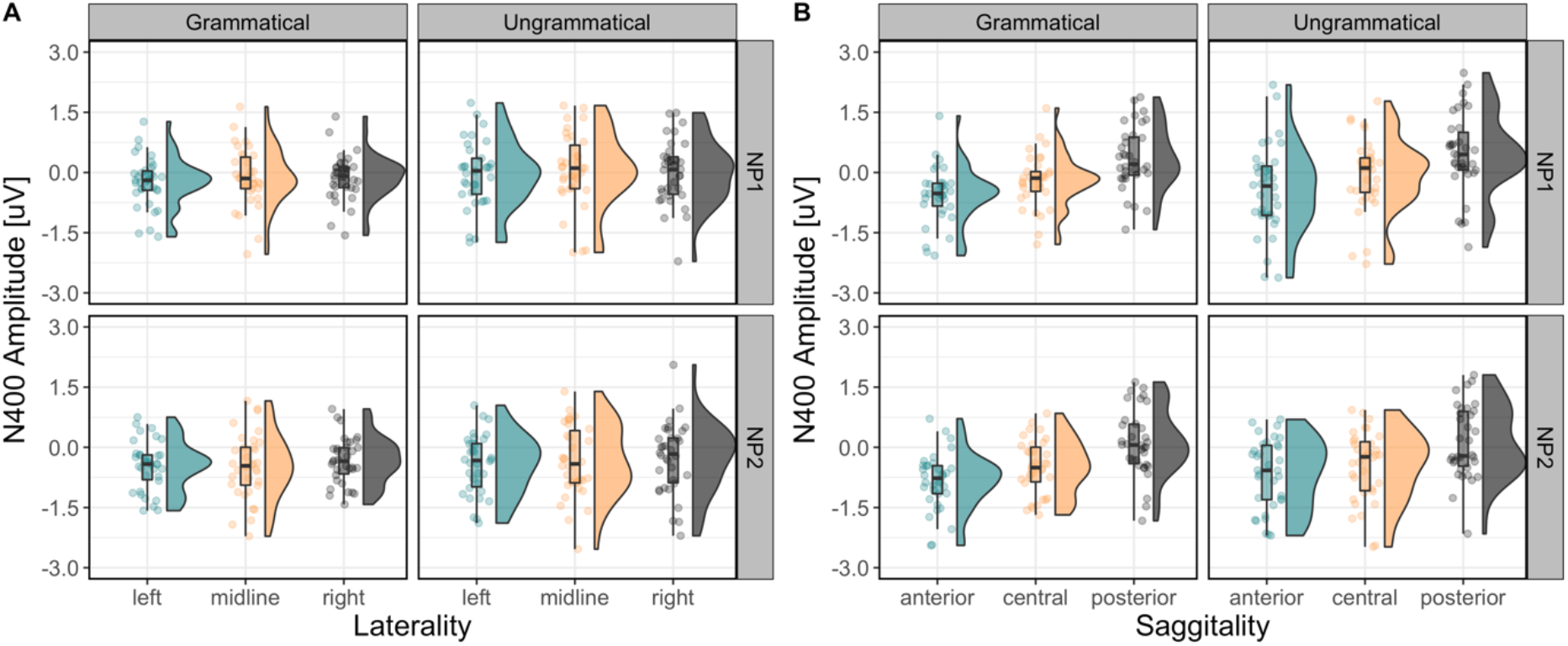
(A) N400 amplitude by grammatical (left) and ungrammatical (right) sentences and first noun phrase (top row) and second noun phrase (bottom row) classifier violations across lateral (A) and sagittal (B) channels. Data points represent single subject N400 estimates.

**Figure 4.**
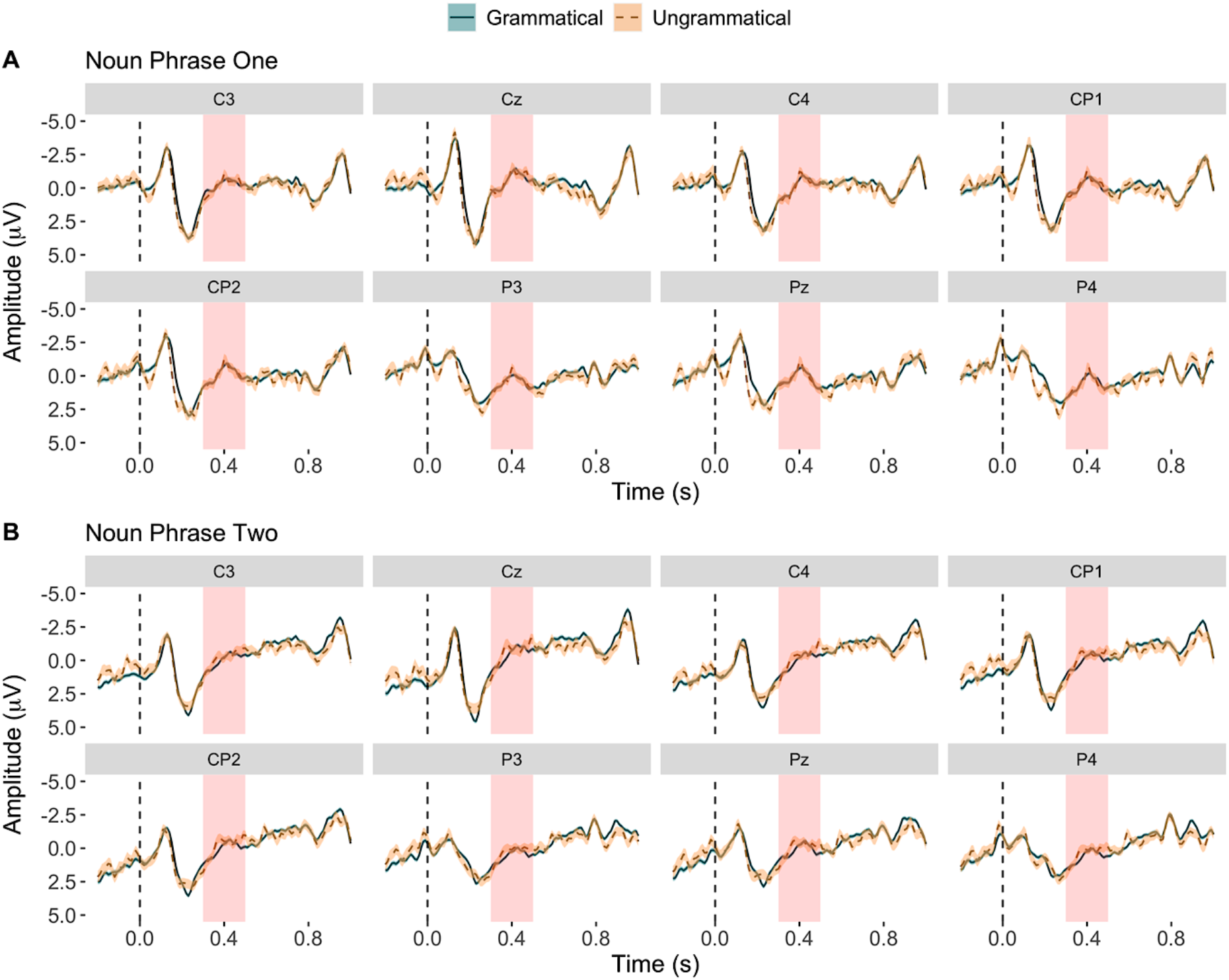
Grand averaged ERPs at the position of violation. The blue solid line represents grammatical sentences, while the orange dashed line represents classifier-noun violations. Amplitude (uV) is represented on the y-axis, while negativity is plotted downwards. Time (in seconds) is represented on the x-axis. The shaded red region represents the time region of the N400 (i.e., 300 – 500 ms).

**Figure 5.**
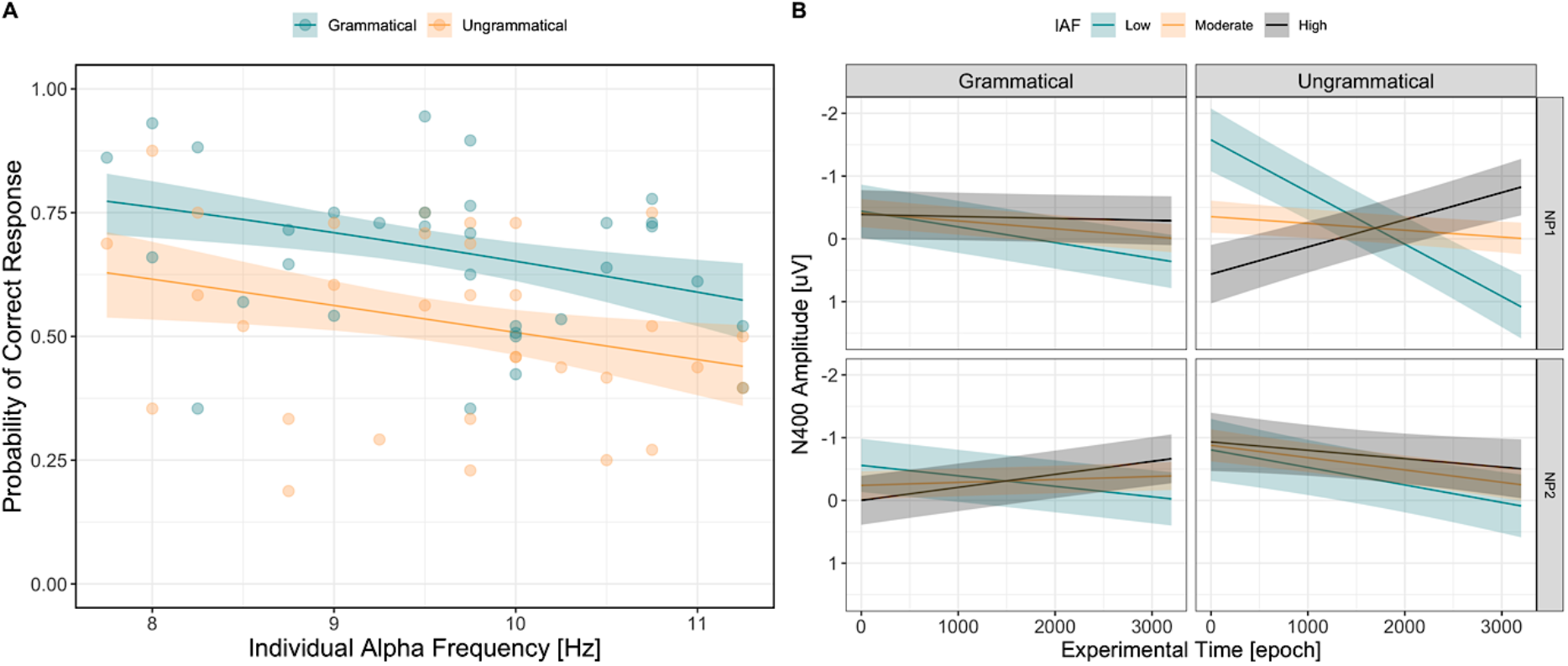
(A) Interaction between IAF and sentence type in predicting the probability of correct response. Data points represent single subject estimates, while the shaded region indicates the 83% confidence interval. (B) Interaction between N400 amplitude (y-axis), IAF (low = blue; moderate = yellow; high = grey), and experimental time (x-axis) for grammatical (left) and ungrammatical (right) classifier-noun pairing violations at the position of the first (top row) and second (bottom row) noun phrase.

### 4.3. Modelling n400 amplitude as a function of time, grammaticality and iaf

To address the remaining hypotheses surrounding the N400, a linear mixed model was constructed. A significant main effect of Grammaticality was found (χ2(1) = 11.09, *p* < .001), indicating that ungrammatical sentences elicited a significantly larger N400 (see Figure 3 and Figure 4 for a visualisation of the distribution of N400 amplitude across lateral and sagittal regions and ERP waveforms, respectively).

A significant interaction between Grammaticality and Epoch was found (χ2(1) = 10.50, *p* = .001), supporting the hypothesis that a significant increase in N400 amplitude at the position of classifier-noun violations over the course of the experiment would be found (for a visualisation of this effect, see Figure 5B). Regarding the hypothesis that N400 amplitude would be stronger for participants with a higher IAF compared to lower, the interaction between grammaticality and IAF was non-significant (χ2(1) = 0.82, *p* = .36). This indicates that N400 amplitude was not significantly more pronounced for participants with a higher IAF compared to those with a lower IAF.

However, a significant four-way Grammaticality × Epoch × IAF × Noun Phrase interaction was found (χ2(1) = 17.15, *p* <.001), partially supporting hypotheses H^5a^ and H^5b^. This interaction is shown in Figure 5B, which illustrates that for classifier-noun violations at the position of the first noun phrase, those with a lower IAF demonstrated a decreased N400 amplitude over the course of the experiment. The inverse relationship is seen in those with a higher IAF, with an increase in N400 amplitude over the experiment; the slope for violations is also steeper than that of grammatical sentences. Finally, those with a moderate IAF estimate demonstrated no change in N400 amplitude over the experiment, with no differences in slope observed between sentence types.

## 5. Discussion

The present study examined both the behavioural and neurophysiological correlates of L2 learning in adulthood and is, to the best of our knowledge, the first to study the relationship between IAF and learning of classifier-noun rules in a complex modified miniature language (MML). H^1^ predicted that behavioural performance would improve over the experiment; however, no significant interaction between experimental time (i.e., block 1 – 6) and grammaticality on the probability of a correct response was found. H^2^ posited that those with a higher IAF would perform better on the grammaticality judgement task than those with a lower IAF; however, this was unsupported as the two-way interaction was significant in the opposite direction – that is, those with a lower IAF performed better at discerning classifier-noun pair violations. Further, a significant four-way interaction was found between IAF, experimental time, grammaticality and noun phrase position on N400 amplitude. These results suggest that a more complex modulating factor underlying adulthood L2 learning than previously hypothesised exists.

### 5.1. Behavioural differences underlying classifier-noun rule learning

A significant effect of grammaticality was observed, demonstrating that participants were better at recognising grammatical than ungrammatical sentences. These findings point to the potentially important role of the learning phase of the experiment in ensuring participants can correctly identify grammatical sentences. However, the variability in individual performance points to the possibility that there are inter-individual differences in learning the grammatical rules underlying classifiers sufficiently to translate into correct grammaticality judgements. Unlike behavioural responses which are given at the end of stimulus presentation, electrophysiological signals measured throughout the experiment provides a plethora of information which can begin to elucidate the neural underpinnings of behavioural judgements. Bornkessel-Schlesewsky and Schlesewsky (2019) hypothesise that negative ERPs, such as the N400, are electrophysiological representations of precision-weighted prediction errors and subsequent model updating in language processing, with errors induced by cues (such as classifier noun-pairings) that are relevant for sentence interpretation having a greater effect on subsequent model updating. While there appears to be a significant, *albeit* small, element of surprise as indexed by the N400, participants seem to be only able to consolidate this learning with varying success over the course of the experimental task for sufficient learning to manifest behaviourally. From this perspective, the large task-evolving effects on the N400 in response to classifier-noun violations likely reflected mechanisms of predictive processing and associated model updating, but these effects were not strong enough to manifest behaviourally.

These findings may further be due to participants being presented with an extensive vocabulary and underlying rules that needed to be learnt over a relatively short period. Although the focus of this study was classifiers, sentence stimuli were part of a broader study involving further grammatical manipulations in Mini-Pinyin regarding word order rules (Cross et al., 2020a; also see Cross et al. 2021). For monolingual native English speakers, word order rules might be more salient than the more ‘arbitrary’ rules governing novel classifier-noun pairings. Word order is more salient in English (MacWhinney et al., 1984) than Mandarin (which has a more flexible word order; Bornkessel-Schlesewsky, 2011; Yao, 2018). As participants were monolingual native English speakers, participants may have been biased towards learning word order rules as a cue for sentence processing over classifier rules (Cross et al., 2020a).

It is also important to note that sentences in Mini-Pinyin are constructed in that the first classifier noun-pair phrase (NP1) precedes the second classifier noun-pair phrase (NP2). This, when looked at with the number of other features comprising Mini-Pinyin, may yet again mean that participants, as the sentence unfolds, are overlooking classifier-noun pair violations in favour of other features which they were biased to seeing as more important in sentence processing (Cross et al., 2020a). These results further support the idea that the degree of N400 amplitude is representative of prediction errors, as participants on the sentence-level are adjusting to the anomality of the classifier-noun pair violations, which may require deeper learning to manifest behaviourally. Future research may warrant the inclusion of a control group of participants whose native language, like Mandarin, contains classifier-like rules (e.g., Japanese), allowing for comparisons between the two to see if native-like ERP components are found, with IAF as a moderating factor.

### 5.2. IAF as a marker of inter-individual differences in classifier-noun rule learning

It is apparent that IAF as a proxy for general cognitive ability modulates the learning of classifier-noun rules but seems to do so in a way contradictory to what was hypothesised. Despite previous research, the hypothesis that a higher IAF would be associated with greater ease in recognising and integrating novel aspects of an MML was not supported. Several possible explanations can potentially elucidate why the inverse of our hypothesis was observed.

Research attempts to link a higher IAF with faster information processing cycles and subsequent greater cognitive ability, specifically *g*-factor intelligence (Grandy et al., 2013; Osaka et al., 1999). However, results from the current study are not in line with this theory. Previous studies have highlighted the possible role alpha rhythms play in processing input in ‘cycles’ of sensory integration (e.g., Mierau et al., 2017). IAF may represent these processing cycles, such that the alpha rhythm reflects the speed at which information is processed within thalamo-cortical and cortico-cortical networks (Kurthen et al., 2020; Surwillo, 1963). By contrast, those with a higher IAF performed significantly worse at making correct grammaticality judgements than those with a lower IAF. It is possible that processing information in faster processing cycles meant that high IAF individuals were less adept at extracting critical information and making a correct judgement during the temporal unfolding of sentential information. The inverse is seen in those with a lower IAF. Instead, presented sentences were likely processed through much slower ‘windows’ of information processing. Previous research (e.g., Howard et al., 2017) looking at the influence of IAF speed on performance on a visual information processing task found that a slower IAF was predictive of superior performance. From this perspective, a slower IAF may reflect attentional processing rather than working memory, meaning low IAF individuals could better attend to the information needing to be processed, and integrate lower level linguistic input (i.e., single words) into larger units (i.e., classifier-noun pairings) in order to generate correct behavioural responses. We also speculate that there is a possibility that grammaticality judgements required attending to and integrating information over a longer time scale, hence why low IAF individuals performed significantly better than those with a high IAF.

Further research can consider this relationship in greater detail to determine whether these findings result from processing cycle speed or if a more complex factor, such as different processing strategies in high vs. low IAF individuals, is underlying the modulatory effect of IAF on classifier rule learning. Nonetheless, this research contributes to the debate surrounding the idea that a ‘faster’ brain is not necessarily advantageous, adding to existing theories on individual differences in information processing (e.g., Hofmans & Mullet, 2011).

### 5.3. The N400, IAF and classifier-noun rule learning across time

We found partial support for our fifth hypothesis, namely that N400 amplitude for classifier violations compared to correct pairings would decrease throughout the experiment and this effect will be larger for high versus low IAF individuals. While we found that N400 amplitude was larger for the first relative to second classifier-noun violation, and that this effect diminished across the experiment, this effect was larger for low relative to high IAF individuals, suggesting that the relationship between IAF, epoch (i.e., experimental time) and classifier-noun violations on N400 amplitude may be more complex than hypothesised.

One explanation is that variability across classifier types may have influenced learning. As previously mentioned, Mini-Pinyin contains four classifier types with corresponding nouns (for a complete list, refer to Table 1). Participants may have learned some classifier-noun pairings better than others so that there is no resultant aggregate effect when collapsing all possible classifier-noun rules. Further, item variability has been highlighted as affecting N400 amplitude, with specific factors such as lexical association modulating N400 amplitude (Kyriaki et al., 2020; Laszlo & Federmeier, 2011). The relatively small number of trials (and by extension, items) may have resulted in relatively high variation in item-related effects on the N400 and behavioural accuracy.

A more theoretically driven account relates to IAF reflecting temporal receptive windows of information processing (Bornkessel-Schlesewsky et al., 2015; Cecere et al., 2015; Samaha & Postle, 2015) and the N400 indexing the locus of prediction error and model updating (Bornkessel-Schlesewsky & Schlesewsky, 2019). From this perspective, individuals with a lower IAF would have had more time to integrate incoming sensory information, likely resulting in stronger internal predictive models, reflected in a reduction in N400 amplitude over time. The potential interaction between IAF, internal model adaptation and time is illustrated in Figure 6.

**Figure 6.**
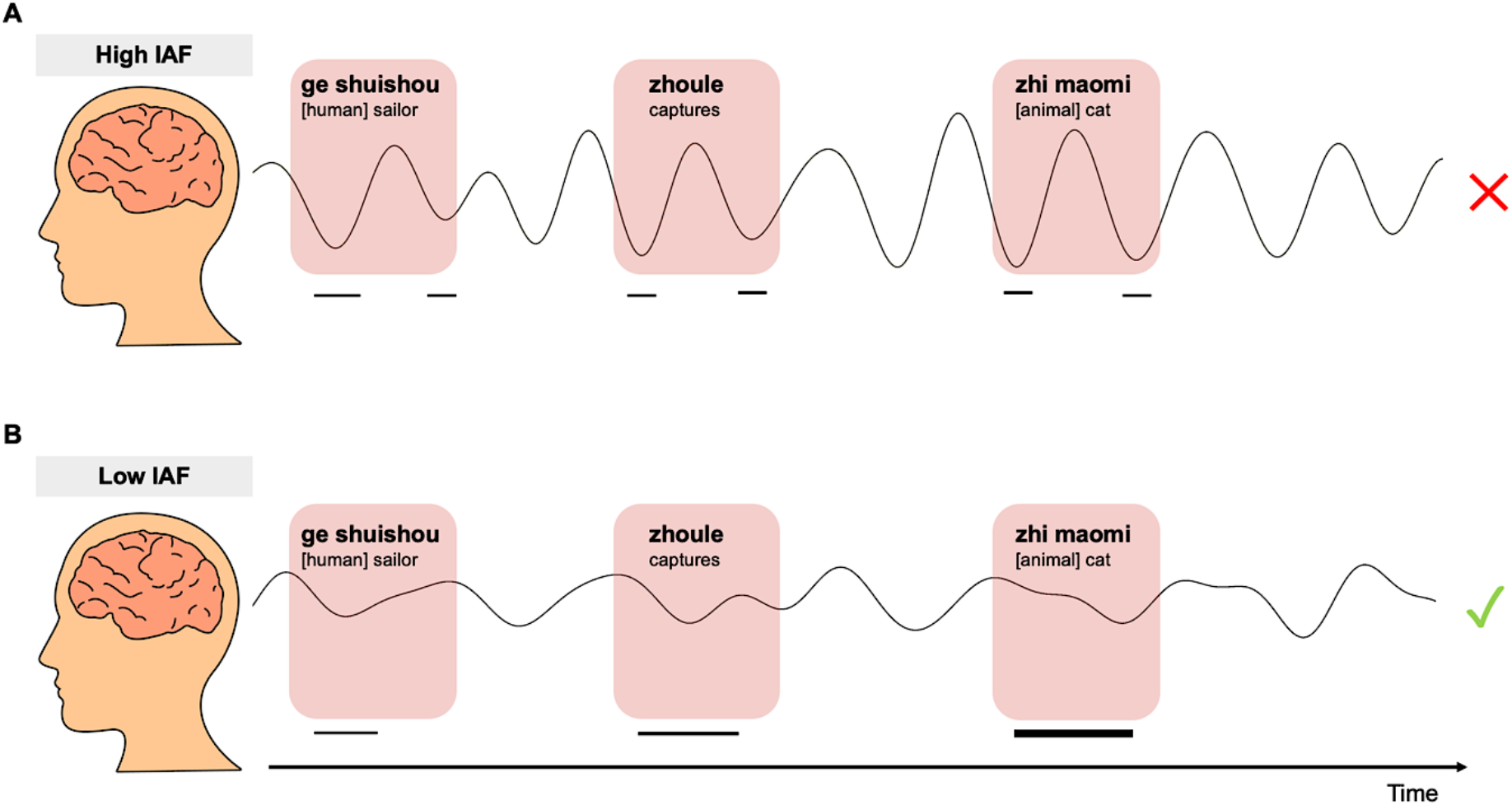
Schematic of the proposed relationship between IAF, internal model adaption and experimental time. (A) High IAF individuals have faster cycles of information processing, resulting in multiple sinusoids during both classifier-noun and verb presentation. (B) Low IAF individuals, with longer temporal receptive windows, are able to uptake incoming information more efficiently, resulting in a strengthening of internal model representations of the linguistic rules, indicated by the systematic thickening of the horizontal black line across time.

### 5.4. Directions for future research

Although previous literature has linked classifier-noun violations to an N400 response, an increasing number of classifier studies have observed left anterior negativity effects (Muller, 2007) and P600 responses (Chan, 2019). Although studies such as Zhang et al. (2012) found that incongruent classifier choice elicited an N400 response, the P600 was also found when the animacy of a classifier-noun pair did not match. Future research necessitates the inclusion of other ERPs in analysis to see if they interact with IAF in a meaningful manner.

Finally, adding to emerging evidence from other domains, further research could also observe how other modulating factors in conjunction with IAF, such as statistical learning ability (SLA), affect language learning. Hauser et al. (2002) pointed to the possibility that the ability to detect regularities in language may have evolved from other domains involving statistical learning. Previous research using artificial grammars has linked SLA to one’s ability to infer abstract, linguistic structures in language learning (Cross et al., 2020a, 2020b; Saffran, 2001), linking variation in SLA with individual L2 learning capabilities (Arciuli & Torkildsen, 2012; Erickson & Thiessen, 2015). Further research warrants looking at how SLA, together with IAF, to determine whether these factors interact to predict language learning.

## 6. Conclusions

This study addresses the gap in previous literature surrounding the role of IAF in adulthood L2 acquisition and is advantaged by its use of an MML, Mini Pinyin, an ecologically valid higher-order language learning paradigm. This study further highlights that possessing a higher IAF is not advantageous in all domains and that the pervasive idea that a high IAF is synonymous with more efficient information processing may not generalize to complex language-related rules. Those with a higher IAF, contrarily, performed worse than low-IAF counterparts. These findings add to the language learning literature by highlighting that more complex processes underlie one’s ability to identify and extract regularities from information input, such as in L2 learning. Theoretically, this study also contributes to the growing neuropsycholinguistic literature surrounding MMLs as a viable method for studying higher-order language learning. Taken together, this study serves as a starting point from which future research can examine the role that intrinsic neural activity, such as the IAF, plays in language learning.

## Acknowledgement

Preparation of this work was supported by Australian Commonwealth Government funding awarded to ZRC under the Research Training Program. IB-S is supported by an Australian Research Council Future Fellowship (FT160100437). We thank Isabella Sharrad, Lena Zou-Williams, Erica Wilkinson, Nicole Vass and Angela Osborn for help with data collection. Thank you also to the participants.

