## supplementary material for "Electrophysiological indices of individual differences in adult language learning"

Table S1: Summary of model produced by the call `glmer(formula = Correct ~ gram * Block + (1 + gram | subj) + (1 | Item_Nm), data = classifiers, family = "binomial", control = glmerControl(optimizer = "bobyqa", calc.derivs = TRUE))`
Generalized linear mixed model fit by maximum likelihood (Laplace Approximation)

| AIC | BIC | logLik | deviance | df.resid |
|---|---|---|---|---|
| 8185 | 8294 | -4077 | 8153 | 6704 |

Scaled residuals:

| Min | 1Q | Median | 3Q | Max |
|---|---|---|---|---|
| -3.79 | -0.94 | 0.5 | 0.73 | 2.75 |

Random effects:

| Groups | Term | Std.Dev. | Corr |
|---|---|---|---|
| Item_Nm | (Intercept) | 0.037036 | |
| subj | (Intercept) | 0.647049 | |
| | gram[Grammatical] | 0.502612 | -0.058 |

Number of obs: 6720, groups: Item$_{N}m, 36; subj, 35$.

Fixed effects:

| | Estimate | Std. Error | z value | Pr(>\|z\|) | |
|---|---|---|---|---|---|
| (Intercept) | 0.38 | 0.11 | 3.3 | 0.00098 | *** |
| gram[Grammatical] | 0.39 | 0.091 | 4.3 | 1.9e−05 | *** |
| Block.L | 0.0027 | 0.077 | 0.035 | 0.97 | |
| Block.Q | −0.05 | 0.076 | −0.66 | 0.51 | |
| Block.C | −0.075 | 0.076 | −0.99 | 0.32 | |
| Block$^4$ | −0.02 | 0.076 | −0.26 | 0.79 | |
| Block$^5$ | −0.13 | 0.075 | −1.7 | 0.095 | . |
| gram[Grammatical]:Block.L | −0.097 | 0.077 | −1.3 | 0.21 | |
| gram[Grammatical]:Block.Q | 0.12 | 0.076 | 1.5 | 0.13 | |
| gram[Grammatical]:Block.C | 0.083 | 0.076 | 1.1 | 0.28 | |
| gram[Grammatical]:Block$^4$ | −0.14 | 0.076 | −1.8 | 0.067 | . |
| gram[Grammatical]:Block$^5$ | 0.11 | 0.075 | 1.5 | 0.13 | |

Table S2: Summary of model produced by the call `glmer(formula = Correct ~ gram * iaf + (1 | subj) + (1 | Item_Nm), data = classifiers, family = "binomial", control = glmerControl(optimizer = "bobyqa", calc.derivs = FALSE))`
Generalized linear mixed model fit by maximum likelihood (Laplace Approximation)

| AIC | BIC | logLik | deviance | df.resid |
|---|---|---|---|---|
| 7634 | 7675 | -3811 | 7622 | 6138 |

Scaled residuals:

| Min | 1Q | Median | 3Q | Max |
|---|---|---|---|---|
| -3.35 | -1.01 | 0.52 | 0.79 | 1.51 |

Random effects:

| Groups | Term | Std.Dev. |
|---|---|---|
| Item_Nm | (Intercept) | 0.02676 |
| subj | (Intercept) | 0.63402 |

Number of obs: 6144, groups: Item$_Nm, 36; subj, 32$.

Fixed effects:

| | Estimate | Std. Error | z value | Pr(>\|z\|) | |
|---|---|---|---|---|---|
| (Intercept) | 2.8 | 1.2 | 2.4 | 0.018 | * |
| gram[Grammatical] | 0.53 | 0.32 | 1.6 | 0.099 | . |
| iaf | −0.24 | 0.12 | −2 | 0.045 | * |
| gram[Grammatical]:iaf | −0.023 | 0.033 | −0.7 | 0.48 | |

Table S3: Summary of model produced by the call `lmer(formula = mean ~ scale(epoch) * iaf * gram * np * prestim + (1 | channel) + (1 | subj) + (1 | item), data = eeg_final_model)`
Linear mixed model fit by REML. t-tests use Satterthwaite's method

REML criterion at convergence: 761263

Scaled residuals:

| Min | 1Q | Median | 3Q | Max |
|---|---|---|---|---|
| -5.97 | -0.63 | -0.01 | 0.62 | 8 |

Random effects:

| Groups | Term | Std.Dev. |
|---|---|---|
| item | (Intercept) | 0.17291 |
| subj | (Intercept) | 0.74185 |
| channel | (Intercept) | 0.23314 |
| Residual | | 4.32875 |

Number of obs: 131888, groups: item, 36; subj, 32; channel, 8.

Fixed effects:

| | Estimate | Std. Error | df | t value | Pr(>\|t\|) | |
|---|---|---|---|---|---|---|
| (Intercept) | 0.16 | 1.3 | 31 | 0.12 | 0.91 | |
| scale(epoch) | 1.4 | 0.19 | 1.3e+05 | 7.3 | 2.1e−13 | *** |
| iaf | −0.043 | 0.14 | 31 | −0.31 | 0.76 | |
| gram[G] | 0.1 | 0.18 | 1.3e+05 | 0.59 | 0.56 | |
| np[N1] | −0.036 | 0.18 | 1.3e+05 | −0.2 | 0.84 | |
| prestim | −0.22 | 0.025 | 1.3e+05 | −8.6 | 6.5e−18 | *** |
| scale(epoch):iaf | −0.13 | 0.02 | 1.3e+05 | −6.8 | 1.2e−11 | *** |
| scale(epoch):gram[G] | −0.58 | 0.19 | 1.3e+05 | −3.1 | 0.002 | ** |
| iaf:gram[G] | −0.0053 | 0.019 | 1.3e+05 | −0.28 | 0.78 | |
| scale(epoch):np[N1] | 0.63 | 0.19 | 1.3e+05 | 3.3 | 0.00082 | *** |
| iaf:np[N1] | 0.016 | 0.019 | 1.3e+05 | 0.86 | 0.39 | |
| gram[G]:np[N1] | 0.49 | 0.18 | 1.3e+05 | 2.7 | 0.0065 | ** |
| scale(epoch):prestim | −0.059 | 0.026 | 1.3e+05 | −2.2 | 0.025 | * |
| iaf:prestim | 0.0032 | 0.0026 | 1.3e+05 | 1.2 | 0.22 | |
| gram[G]:prestim | 0.05 | 0.025 | 1.3e+05 | 2 | 0.046 | * |
| np[N1]:prestim | 0.022 | 0.025 | 1.3e+05 | 0.87 | 0.38 | |
| scale(epoch):iaf:gram[G] | 0.055 | 0.02 | 1.3e+05 | 2.8 | 0.0056 | ** |
| scale(epoch):iaf:np[N1] | −0.062 | 0.02 | 1.3e+05 | −3.2 | 0.0015 | ** |
| scale(epoch):gram[G]:np[N1] | −0.74 | 0.19 | 1.3e+05 | −3.9 | 8.9e−05 | *** |
| iaf:gram[G]:np[N1] | −0.057 | 0.019 | 1.3e+05 | −3.1 | 0.0021 | ** |
| scale(epoch):iaf:prestim | 0.007 | 0.0027 | 1.3e+05 | 2.6 | 0.01 | * |
| scale(epoch):gram[G]:prestim | 0.031 | 0.026 | 1.3e+05 | 1.2 | 0.23 | |
| iaf:gram[G]:prestim | −0.0047 | 0.0026 | 1.3e+05 | −1.8 | 0.071 | . |
| scale(epoch):np[N1]:prestim | 0.031 | 0.026 | 1.3e+05 | 1.2 | 0.23 | |
| iaf:np[N1]:prestim | −0.0019 | 0.0026 | 1.3e+05 | −0.73 | 0.47 | |
| gram[G]:np[N1]:prestim | −0.15 | 0.025 | 1.3e+05 | −6.1 | 1.1e−09 | *** |
| scale(epoch):iaf:gram[G]:np[N1] | 0.082 | 0.02 | 1.3e+05 | 4.2 | 3.1e−05 | *** |
| scale(epoch):iaf:gram[G]:prestim | −0.0032 | 0.0027 | 1.3e+05 | −1.2 | 0.24 | |
| scale(epoch):iaf:np[N1]:prestim | −0.0036 | 0.0027 | 1.3e+05 | −1.3 | 0.19 | |
| scale(epoch):gram[G]:np[N1]:prestim | −0.064 | 0.026 | 1.3e+05 | −2.5 | 0.014 | * |
| iaf:gram[G]:np[N1]:prestim | 0.016 | 0.0026 | 1.3e+05 | 6.2 | 7.2e−10 | *** |
| scale(epoch):iaf:gram[G]:np[N1]:prestim | 0.0067 | 0.0027 | 1.3e+05 | 2.5 | 0.013 | * |
